# Non-micelle-like Amyloid Aggregate Stabilizes Amyloid β (1-42) Growth Nuclei Formation

**DOI:** 10.1101/2022.12.09.519846

**Authors:** Ikuo Kurisaki, Shigenori Tanaka

**Author notes:** **Corresponding Authors** Ikuo Kurisaki - Waseda Research Institute for Science and Engineering, Waseda University, Tokyo 169-8555, Japan;, Shigenori Tanaka - Graduate School of System Informatics, Kobe University, 1-1 Rokkodai, Nada-ku, Kobe 657-8501, Japan. **Data availability statement** The data that support the findings of this study are available from the corresponding authors upon reasonable request. **Ethics approval statement** None. **Patient consent statement** None. **Permission to reproduce material from other sources** There are no reproduced materials from other sources. All of the materials were newly created for this manuscript. **Clinical trial registration** None.

## Abstract

Protein aggregate formations are essential processes to regulate biochemical networks in the cell, while anomalously formed aggregates such as amyloid fibrils cause serious neuronal diseases. It has been discussed for a quarter century that protein crowding milieus, such as micelle-like aggregates, promote the formation of growth nuclei, fibril-growth competent aggregates which trigger rapid growth of pathogenic amyloid fibrils, but the mechanisms are still elusive, in particular at microscopic level. In this study, we examined the long-standing problem by employing atomistic molecular dynamics simulations for amyloid β(1-42) (Aβ_42_), the paradigmatic amyloid-forming peptide. First, we constructed an atomistic model of Aβ_42_ growth nuclei in Aβ_42_ aggregate milieu, the pentameric Aβ_42_ protomer dimer surrounded by 40 Aβ_42_ monomers. Next, we simulated Aβ_42_ monomer dissociation from the Aβ_42_ growth nuclei and examined the effect of Aβ_42_ aggregate milieu on the process. Aβ_42_ aggregates spatially restrict Aβ_42_ monomer dissociation pathways, while such spatial restriction itself does not significantly suppress Aβ_42_ monomer dissociation from the growth nuclei. Rather, Aβ_42_ aggregate milieus thermodynamically stabilize an Aβ_42_ monomer binding to the growth edge by making atomic contacts with the monomer and contributes to stable formation of growth nuclei.

A part of the aggregate milieu anchors dissociating monomer to the remaining part of growth nuclei, suggesting cooperative suppression of Aβ_42_ monomer dissociation from Aβ_42_ growth nuclei. Since the Aβ_42_ aggregate milieu does not take a micelle-like configuration, we here discuss a new mechanism for stable formation of Aβ_42_ growth nuclei in the presence of aggregate milieu.

## Introduction

Experimental studies performed within the last decades remarkably updated our understanding for biological importance of aggregation formation of biological molecules. Critical biological events (*e*.*g*., the gene transcription, protein synthesis, and signal transduction) progress through the aggregate formation and dissolution^1-5^, thus denoting that disorder of the functional protein aggregates can directly lead to serious diseases. Nonetheless, we have not sufficiently understood how such aggregates form and function in the cell, even for the paradigmatic molecular species such as amyloid β protein.

Over the past 40 years, formation mechanisms of protein aggregates and the biological roles have been widely examined with regard to serious neurodegenerative diseases such as Alzheimer and Parkinson diseases^6^. Since anomalous protein aggregates are supposed to be pathogenic causes of these diseases, one representative subject is the understanding of pathogenic amyloid fibril formation^7^. Then, it has been one of the central problems how we can suppress formation of pathogenic amyloid fibrils, with the aim of developing therapeutic strategy for these serious diseases^8,9^.

To understand the aggregate formation process at molecular level, the kinetic models of aggregate formation have been extensively studied and established^10-12^. Amyloid fibril growth starts from accumulation of sufficient amounts of growth nuclei species, which are found among the repertoire of oligomeric assembly of proteins and are classified into fibril-like, growth-competent aggregates^11^. This molecule-level mechanism of amyloid fibril formation indicates that clarifying the minimum size of growth nuclei species is a landmark knowledge to understand physicochemical mechanisms of dramatic progress of pathogenic amyloid formation.

To solve the problem, nano-scale observation techniques have been widely employed to observe oligomerization processes in single molecule-level resolution^13-15^. As a technical counterpart to give microscopic insights, theoretical simulations have been extensively employed and have revealed key atomistic interactions acting on aggregate-prone proteins through their aggregation process.^16^ These studies have mainly focused on amyloid proteins themselves, which work as building blocks of aggregates and form specific interaction acting among fibril-growth competent monomers.

Besides such specific interactions acting among amyloid proteins in fibril-like shapes, it has been suggested that protein crowding milieus, which consist of the same amyloid protein and form non-fibril, micelle-like shape, also contribute to stable formation of growth nuclei, which has been discussed by several experimental studies for the paradigmatic amyloid peptide, amyloid β (1-42) (Aβ_42_) within the past two decades^17-21^.

However, the mechanisms of aggregate crowding milieu for stable formation of growth nuclei have not been sufficiently elucidated, due to technical difficulty to experimentally capture the molecular structure and dynamics. The emergence of such Aβ_42_ aggregate crowding milieu is transient during amyloid aggregates formation process, then often being out of time resolution of experimental observations. Besides, the crowding milieus are essentially heterogeneous ensemble of conformationally disordered Aβ_42_ monomers, thus being challenging to determine the atomic structures of aggregate crowding milieus so far. Recalling these molecular natures of aggregate crowding milieus, we could suppose atomistic simulations as a feasible, computational counterpart to consider the problem.

In the present study, we examine the effects of Aβ_42_ aggregate crowding milieu on Aβ_42_ stable formation of Aβ_42_ growth nuclei and the microscopic origin. We start from testing the speculation, which we made in the previous study^21^: an Aβ_42_ aggregate crowding milieu suppresses Aβ_42_ monomer dissociations from a growth edge of Aβ_42_ growth nuclei and prevents disassembly of the growth nuclei. To overcome the above experimental difficulty, we theoretically modeled the atomic configuration of Aβ_42_ aggregate milieu by employing the hybrid Monte Carlo/Molecular Dynamics (MC/MD) approach, which we had developed to examine dynamic processes of multimeric proteins^22,23^. By using the simulation-derived Aβ_42_ aggregate milieu systems for the free energy calculation of Aβ_42_ monomer dissociation from the protomer dimer, we confirmed the stabilization effect of Aβ_42_ aggregate milieus on Aβ_42_ growth nuclei formation via Aβ_42_ suppressing monomer dissociation from the growth edge.

## Methods

### Setup of amyloid-β (1-42) protomer dimer systems

We used the cryogenic electron microscopy (cryo-EM) structure (PDB entry: 5OQV) ^24^ to construct amyloid-β (1-42), Aβ_42_, 25-mer protomer dimer systems (**Figure 1A**); a protomer denotes an Aβ_42_ oligomer composed of Aβ_42_ monomers in fibril growth-competent conformation. We considered the 50 Aβ_42_ monomers by recalling the earlier studies which reports the number range of Aβ_42_ monomer in the non-fibril aggregates milieu^18-20^. This model is annotated by Aβ_42_(25:25) or simply 25:25, hereafter. We designed Aβ_42_(25:25) in this study, by recalling the earlier experimental studies which reported that a micelle-like Aβ_42_ aggregate consists of 25 to 50 Aβ_42_ monomers^18,19^. Although there is another full-length Aβ_42_ fibril structure (PDB entry: 2NAO^25^), we selected this 5OQV structure by considering the relationship with our earlier studies^21,26^.

**Figure 1.**
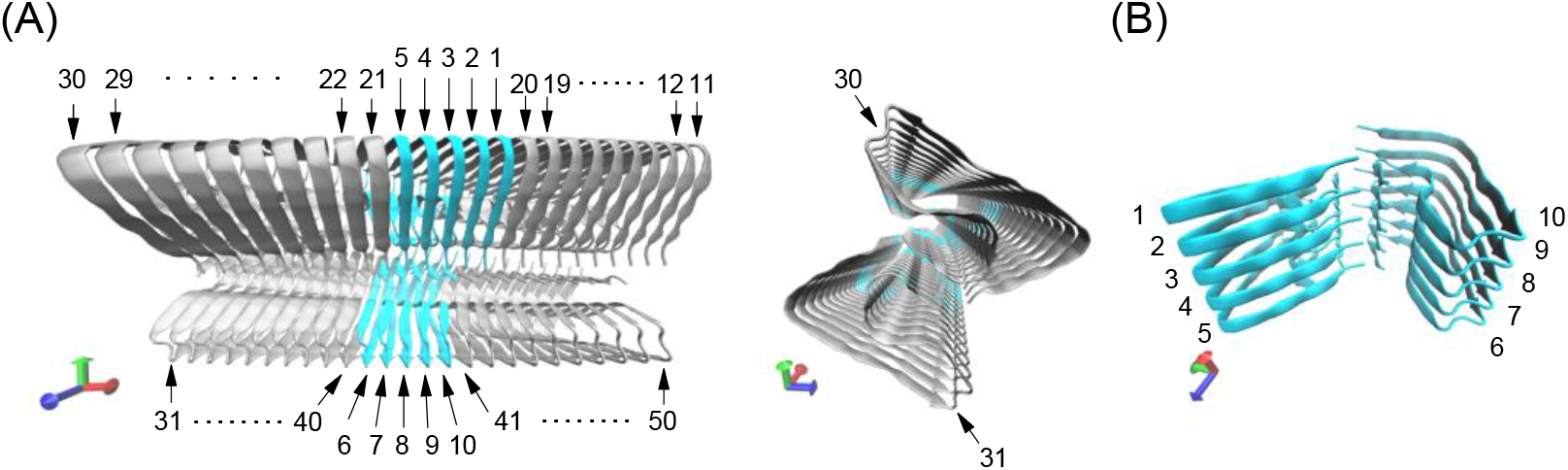
Amyloid β (1-42) 25-mer protomer dimer. (A) the 40 subunits to construct aggregate milieu, and the 5-mer dimers for model of amyloid fibril growth nuclei (referred to as Aβ_42_(5:5)), which are colored with gray and cyan, respectively. Each of the 50 monomers is numbered for the following discussion. (B) Aβ_42_(5:5) whose fibril growth edges are annotated by 1, 5, 6 and 10.

Nε protonation state was employed for each of histidine residues, and all carboxyl groups in aspartate and glutamate residues were set to the deprotonated state. Aβ_42_(25:25) was solvated in the 138 Å×138 Å×180 Å rectangular box, and 97330 water molecules were finally included in the box. Furthermore, 150 K^+^ molecules were added to the solvated system to electrically neutralize the system. Since we are interested in suppression effect of Aβ_42_-crowding milieu on monomer dissociation from Aβ_42_ oligomer (**Figure 1B**), no biological co-solutes were added into aqueous solution except for the counter ions to electronically neutralize these molecular systems. The additional detail for system construction is described in Supporting Information (see **SI-1**).

### Simulation setup

Molecular mechanics (MM) and molecular dynamics (MD) simulations were performed under the periodic boundary condition with GPU-version PMEMD module in AMBER 17 package^27^ based on SPFP algorism^28^ with NVIDIA GeForce GTX1080 Ti. Electrostatic interaction was treated by the Particle Mesh Ewald method, where the real space cutoff was set to 0.9 nm. The vibrational motions associated with hydrogen atoms were frozen by SHAKE algorithm through MD simulations. The translational center-of-mass motion of the whole system was removed by every 500 steps to keep the whole system around the origin, avoiding an overflow of coordinate information from the MD trajectory format. To calculate the forces acting among atoms, AMBER force field 14SB^29^, TIP3P water model^30,31^, and JC ion parameters adjusted for the TIP3P water model^32,33^ were used for amino acid residues, water molecules, and ions, respectively. Molecular modeling of each Aβ_42_(N:N) system was performed using the LEaP modules in AmberTools 17 package^27^. These simulation conditions mentioned above were common in all of the simulations discussed in this manuscript.

### Unbiased MD simulations

Following temperature and density relaxation simulations, 10-ns NPT MD simulations (300 K, 1 bar) were performed and used for following analyses. In the present study, we employed this simulation time length for conformation relaxation of Aβ_42_ protomer dimers via transfer from solid phase to liquid phase.

The system temperature and pressure were regulated with Berendsen thermostat^26^ with a 5-ps of coupling constant and Monte Carlo barostat with attempt of system volume change by every 100 steps, respectively. A set of initial atomic velocities was randomly assigned from the Maxwellian distribution at 0.001 K at the beginning of the NVT simulations. The time step of integration was set to 2 fs. The further details are shown in Supporting Information (see **SI-2**).

### Modeling amyloid aggregate milieu with hybrid MC/MD simulation

The 40 monomers, which are the part of Aβ_42_(25:25), are dissociated to prepare amyloid aggregate milieu by repetitive steered MD simulations. The sequential dissociations of monomers were carried out in the framework of hybrid MC/MD method, which we employed to study disassembly processes of the homomeric protein multimer^22^.

In the first 100 hybrid MC/MD cycles, a set of the four monomers at the fibril growth edges (numbered by 11, 30, 31 and 50 in **Figure 1A**) are considered for dissociation. In the next 100 hybrid MC/MD cycles, another set of four monomers (numbered by 12, 29, 32 and 49), which make contacts with the 11, 30, 31 and 50 monomers, are dissociated. Such a simulation procedure was repeated by changing a set of four monomer (see **Table 1** for simulation setup for each of the hybrid MC/MD simulations) until Aβ_42_(5:5) is left in the aqueous solution. As for the monomers making contacts with Aβ_42_(5:5) (numbered by 20, 21, 40 and 41), we employed 400 hybrid MC/MD cycles. This sequential dissociation procedure is designed due to such a physicochemical intuition that an Aβ_42_ monomer at the growth edge is dissociated from Aβ_42_ growth nuclei more frequently than that in the growth nuclei, which is stacked by two neighboring Aβ_42_ monomer.

**Table 1.** hybrid MC/MD simulation setup for modeling Aβ_42_ aggregate milieu system.

| hcbMC/MD<br>simulation order | number of cycle | <sup>†</sup>monomer pairs undergoing dissociation |
| --- | --- | --- |
| 1 | 100 | 11-12; 30-29; 31-32; 50-49 |
| 2 | 100 | 12-13; 29-28; 32-33; 49-48 |
| 3 | 100 | 13-14; 28-27; 33-34; 48-47 |
| 4 | 100 | 14-15; 27-26; 34-35; 47-46 |
| 5 | 100 | 15-16; 26-25; 35-36; 46-45 |
| 6 | 100 | 16-17; 25-24; 36-37; 45-44 |
| 7 | 100 | 17-18; 24-23; 37-38; 44-43 |
| 8 | 100 | 18-19; 23-22; 38-39; 43-42 |
| 9 | 100 | 19-20; 22-21; 39-40; 42-41 |
| 10 | 400 | 20-1; 21-5; 40-6; 41-10 |

<sup>†</sup> For each monomer pair connected by hyphen, a digit at the left side is assigned to Aβ <sub>42</sub> monomer at the edge of growth axis. A digit is assigned to Aβ <sub>42</sub> monomer as defined in Figure 1.

Inter-Aβ_42_ monomer dissociation was simulated by using steered MD method with force constant of 4.184 kJ/mol/nm^2^. A steered force was set along the distance between the centers of mass of given Aβ_42_ monomer pair. At each hybrid MC/MD cycle, a pair of Aβ_42_ monomers is randomly selected among the candidates shown in **Table 1**. The center of mass (COMs) of Aβ_42_ monomer is defined by using 42 Cα atoms of the Aβ_42_ monomer.

Finally, we obtained the snapshot structure of Aβ_42_(5:5) surrounded by an Aβ_42_ aggregate milieu. This snapshot structure is annotated by Aβ_42_(M), where M denotes a milieu. Using Aβ_42_(M), we performed the 100-ns NPT simulation to relax atomistic interaction between Aβ_42_(5:5) and Aβ_42_ aggregate milieu. The simulation result is annotated by Aβ_42_(R|M), where R denotes the relaxation of the Aβ_42_ with aggregate milieu system.

We also construct Aβ_42_(5:5) isolated in aqueous solution by removing the 40 dissociated Aβ_42_ monomers and 120 K^+^ ions; this snapshot structure is annotated by Aβ_42_(P), where P denotes pure water. The system density was equilibrated by 10 ns-NPT MD simulation. During the above simulation, we fix Aβ_42_(5:5) conformation around that in Aβ_42_(M) with the aim of fairly evaluating the effect of presence of Aβ_42_ aggregate milieus, where harmonic positional restraints with force constant of 20.92 kJ/mol/nm^2^ are imposed on the main chain atoms of Aβ_42_ monomers. Since the simulation setup to obtain Aβ_42_(R|M) and Aβ_42_(P) is similar to that for the above 10-ns NPT MD simulation, we do not repeat the explanation here. Further technical details of the hybrid MC/MD simulations and the procedure to prepare Aβ_42_(P) are given in Supporting Information (see **SI-3** and **SI-4**).

### Umbrella sampling MD simulation combined with steered MD simulation

The dissociation processes of Aβ_42_ monomer from the fibril growth edge were described by combining a steered molecular dynamics (SMD) simulation with umbrella sampling molecular dynamics (USMD) simulations. We defined the reaction coordinate as inter-COM distance between the dissociation monomer (S10) and the remaining of the Aβ_42_ pentamer (S6, S7, S8 and S9).

An SMD simulation was employed to obtain an atomic trajectory of Aβ_42_ monomer dissociation from the remaining part of the Aβ_42_ pentamer. 1-ns SMD simulation was carried out under constant NPT condition (300 K, 1 bar), where the system temperature and pressure were regulated by Langevin thermostat with 1-ps^-1^ collision coefficient, and Berendsen barostat^34^ with a 5-ps coupling constant, respectively. The value of reaction coordinate was gradually changed through the SMD simulations by imposing the harmonic potential with the force constant of 4.184 kJ/mol/nm^2^. The remaining part of Aβ_42_(5:5) was restrained around the initial atomic coordinates by imposing harmonic potential with force constant of 4.184 kJ/mol/nm^2^ on the heavy atoms in the main chains.

Then, certain numbers of snapshot structures were extracted from the SMD trajectory with 0.1 nm interval along the reaction coordinate and employed for USMD windows (shown in **Tables S1, S2** and **S3**). Following temperature relaxation simulations, several nanoseconds-length NVT USMD simulations (300 K) were performed for each of the USMD windows. The system temperature was regulated using Langevin thermostat with 1-ps^-1^ collision coefficient. Each of the last 3-ns USMD trajectories was used to construct a potential of mean force (Convergence of each PMF profile is discussed in **Fig. S1**).

We also performed SMD-combined USMD simulations for an Aβ_42_ monomer, which is a part of amyloid aggregate milieu, S19. The reaction coordinate is the distance between COMs of the monomer and the Aβ_42_ pentamer containing S10. The setup of USMD windows is discussed in **Tables S4** and **S5**. The other simulation conditions are similar to those for the dissociating simulations of S10.

The further details, such as setup of harmonic potentials for each of USMD simulations, are described in Supporting Information (see **SI-5**). The reasons for selecting S10 and S19 will be explained in Results and Discussion section.

### Trajectory analyses

Root mean square deviation (RMSd), hydrogen bond (HB), solvent accessible surface area (SASA) and secondary structure annotation based on DSSP^28^ were calculated with the cpptraj module in AmberTools 17 package^27^. We calculated RMSd to the Aβ_42_ protomer dimer structure derived from the cryo-EM model^21,24^ using the backbone heavy atoms (i.e., C_α_, N, C and O). The geometrical criterion of HB formation is as follows: H-X distance was < 0.35 nm and X-H-Y angle was > 120°, where X, Y and H denote acceptor, donor and hydrogen atoms, respectively.

Buried SASA is defined by the following formula, SASA(A:B) − SASA(A) − SASA(B), where A and B are a molecule or molecular assembly, and A:B denotes the complex of A and B. Besides, the hydrodynamic radius (*R*_*hyd*_), which is defined as **Eq. 1** was used to structurally characterize the hMC/MD simulation-derived aggregates.

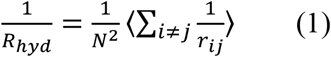

*r*_*ij*_ and *N* denote the distance between the centers of mass of Aβ_42_ monomers composing an Aβ_42_ aggregate (indexed by *i* and *j*) and the total number of such Aβ_42_ monomers, respectively.

Each set of USMD trajectories was used to calculate potential of mean force (PMF) with Weighed Histogram Analysis Method (WHAM)^35^. Statistical errors of PMF values, *σ*_*PMF*_(*ξ*), were estimated by employing bootstrapped sampling^36^:

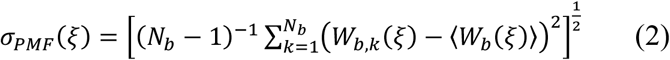

Here, *N*_*b*_, *ξ*, and *W*_*b,k*_ (*ξ*) denote the number of bootstrap sampling, the reaction coordinate and the value of *k*^th^ bootstrapped potential of mean force at each point of *ξ*, respectively. *W*_*b*_ (*ξ*) is average over all *W*_*b,k*_ (*ξ*), where the value of *N*_*b*_ is set to 200 according to the previous study^36^.

We evaluated spatial extension of Aβ_42_ monomer dissociating trajectories by using the bundle radius, *R*_*t*_.

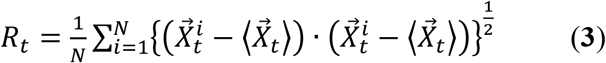

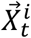 is the center of mass coordinate of dissociating monomer, which is calculated from a set of C_α_ atoms. The superscript, *i* and subscript, *t*, denote a simulation index, which ranges from 1 to N, and the time point in the SMD simulation, respectively. The number of SMD simulation, N, is set to 50 here. The angle parentheses denote average-calculating operation at the time point, *t*, over the *N* independent SMD simulations.

Molecular structures were illustrated using Visual Molecular Dynamics (VMD)^37^. Error bars are calculated from standard error and indicate 95% confidence interval if there is no annotation.

## Results and Discussion

### Simulation-derived Aβ_42_ aggregate milieu consists of fibril-growth incompetent monomers and makes contacts with amyloid pentamer dimer on the fibril growth edge

We generated Aβ_42_ aggregate milieu around Aβ_42_(5:5) by dissociating 40 monomers from Aβ_42_(25:25) with the hybrid MC/MD method as follows. First, we performed 10-ns NPT MD simulation to relax the Aβ_42_(25:25) structure upon transfer from solid phase to aqueous phase. Within the 10-ns NPT MD simulation, both the Aβ_42_(25:25) appear to be relaxed in terms of RMSd convergence (**Figure 2**). Similarly, we found RMSd convergence for the Aβ_42_(5:5), which is positioned in the middle of the Aβ_42_(25:25) (colored by blue in **Figure 1**) and is employed as a model of the growth nuclei in this study.

**Figure 2.**
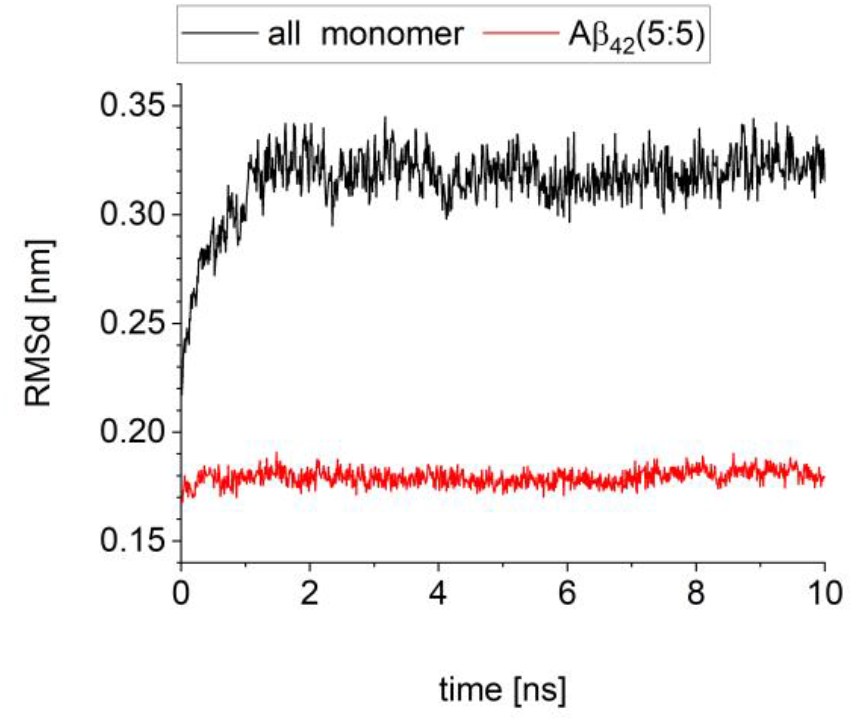
Time course change of root mean square deviation to the experimentally resolved structure. (A) whole of the protofibril, Aβ_42_(25:25) (black line). (B) Pentamer dimer, Aβ_42_(5:5) which is located at the center of the protofibril and colored by blue in **Figure 1** (red line).

Then, we performed the hybrid MC/MD simulations for the10-ns NPT MD snapshot structure and obtained a set of atomic coordinates of Aβ_42_ growth nuclei surrounded by Aβ_42_ aggregate milieu (**Figure 3A**). We annotate the hybrid MC/MD simulation-derived snapshot structure as Aβ_42_(M) hereafter (**Figure 3A**), where the M in the parentheses denotes a ‘milieu’.

**Figure 3.**
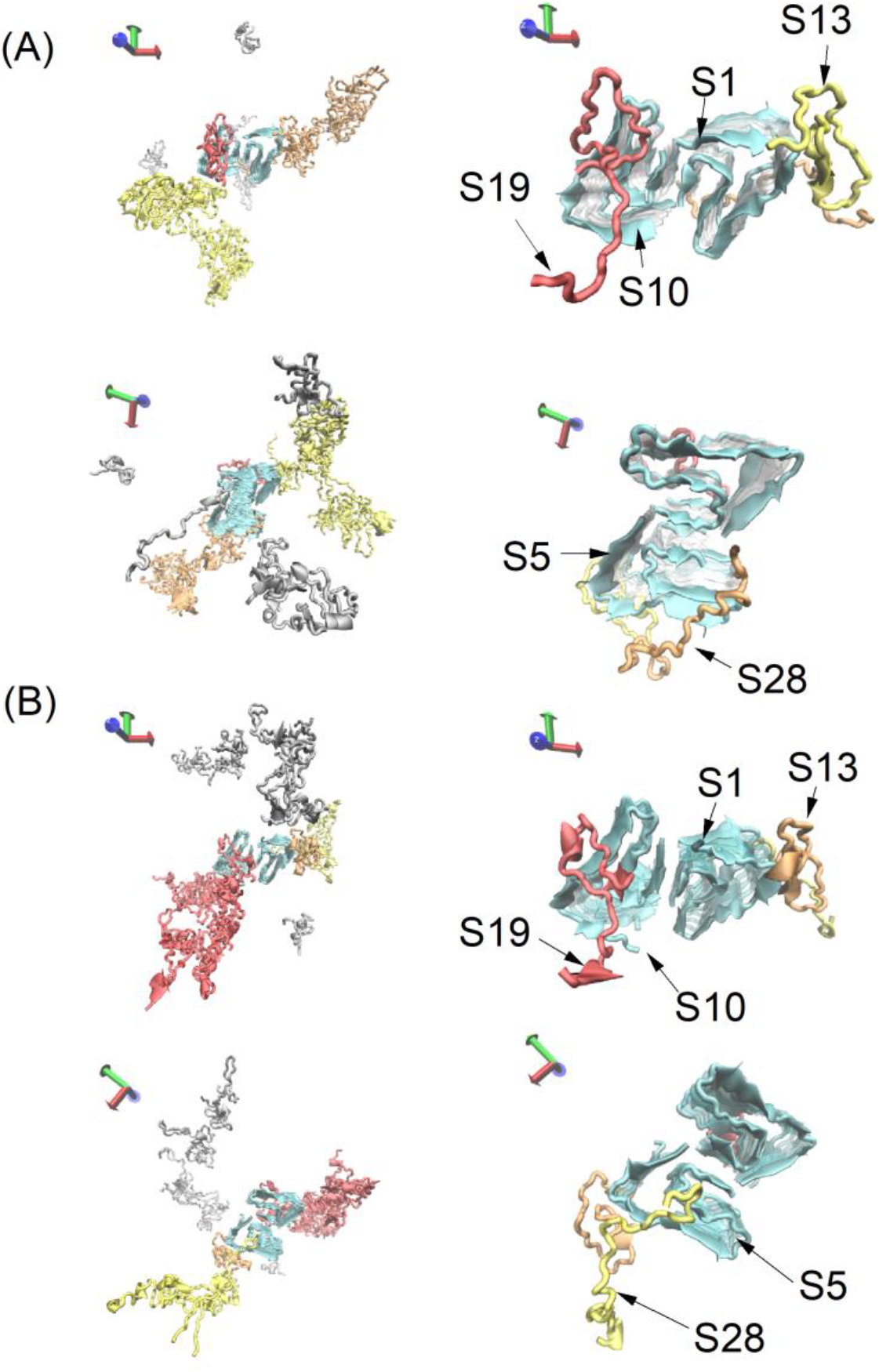
The amyloid β (1-42) (Aβ _42_) pentamer dimer (a model of fibril growth nuclei, Aβ_42_(5:5)) surrounded by the unstructured Aβ_42_ aggregates consisting of 40 monomers. (A) Aβ_42_(M). (B) Aβ _42_(R|M). Aβ_42_(5:5) is colored by cyan. The Aβ_42_ aggregates interacting with Aβ_42_(5:5) are colored by red, yellow and orange. In each of the panels, the whole picture is given in the left, while the monomers which makes contacts with the growth edge of Aβ_42_(5:5) are highlighted in the right.

As remarked in Materials and Methods section, we used Aβ_42_(M) to generate a snapshot structure of Aβ_42_(5:5) in a pure water. We removed a set of atomic coordinates for the 40 Aβ_42_ monomers (indexed from 11 to 50 in **Figure 1A**), which forms aggregate milieu in Aβ_42_(M). This snapshot structure is referred to as Aβ_42_(P), where it is noted that the letter ‘P’ in the parentheses comes from *pure* water. We can consider this system as a reference to discuss the effect of Aβ_42_ aggregate milieu on suppressing Aβ_42_ monomer dissociation from the fibril growth edge.

We also performed a 100-ns NPT MD simulation for Aβ_42_(M) to relax the interaction between the Aβ_42_ growth nuclei and the Aβ_42_ aggregate milieu. The snapshot structure resulted from the simulation is referred to as Aβ_42_(R|M) (**Figure 3B**), where the letter ‘R|M’ in the parentheses denotes the relaxation of Aβ_42_(M). Since the number of hydrogen bond (H-bond) formation between Aβ_42_ growth nuclei and Aβ_42_ aggregate milieu is assumed to reach convergence by 20 ns (see **Figure 4**), we use the 100-ns NPT MD snapshot structure to examine the effect of atomistic interaction between Aβ_42_ growth nuclei and Aβ_42_ aggregate milieu on suppressing Aβ_42_ monomer dissociation from the fibril growth edge.

**Figure 4.**
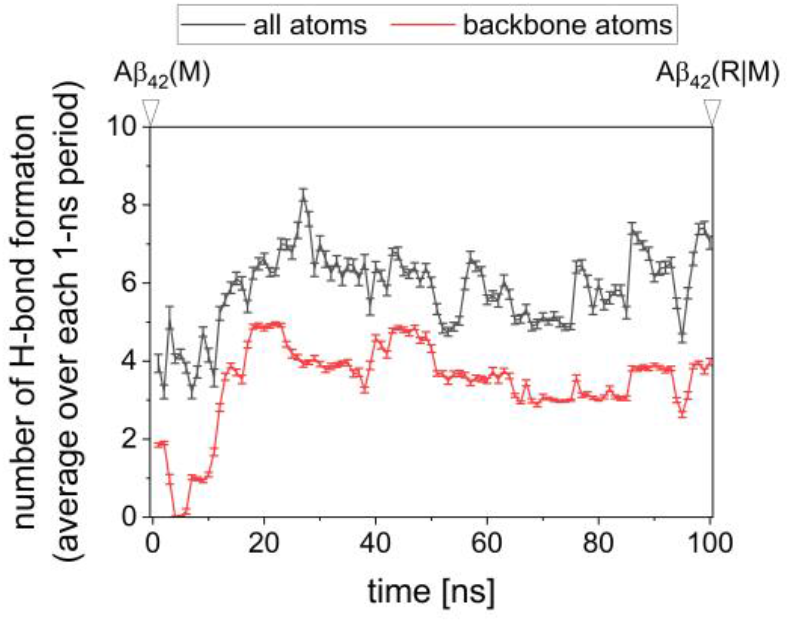
Hydrogen bond (H-bond) formation between Aβ_42_ growth nuclei (the pentamer dimer) and Aβ_42_ aggregate milieu (the remaining Aβ_42_ 40 monomers). The black line denotes H-bond calculated for all atoms, while the red line denotes that for backbone atoms. The H-bond number is averaged over each 1-ns period and the error bars indicate 95% confidence interval.

For both of the Aβ_42_(M) and Aβ_42_(R|M), the components of the Aβ_42_ aggregate milieu appear to be off-pathway, fibril-growth incompetent oligomers (**Figure 3A** and **Figure 3B**): they are probably deviated from the amyloid fibril-growth competent form, that is, a LS shape conformation, which is the building block of the Aβ_42_ fibril structure registered with PDB entry: 5OQV^24^. To confirm this observation, we analyzed the structural properties of Aβ_42_(5:5) and Aβ_42_ aggregate milieu consisting of 40 Aβ_42_ monomers by using the two snapshot structures of Aβ_42_(M) and Aβ_42_(R|M). It is noted that Aβ_42_(P) is obtained from Aβ_42_(M), thus denoting that the two snapshot structures are enough to discuss structural properties of Aβ_42_ monomer.

We examined secondary structure (SS) annotation for each of 50 Aβ_42_ monomers. The monomers in the Aβ_42_(5:5), indexed from 1 to 10 (*discussed in* **Figure 1**), basically take turn or bent structure around 10^th^, 25^th^ and 37^th^ residues (see **Figure 5A** and **5B**), indicating that each of these 10 monomers takes the LS shape. The remaining part of the Aβ_42_(5:5) does not show any characteristic SS motifs.

**Figure 5.**
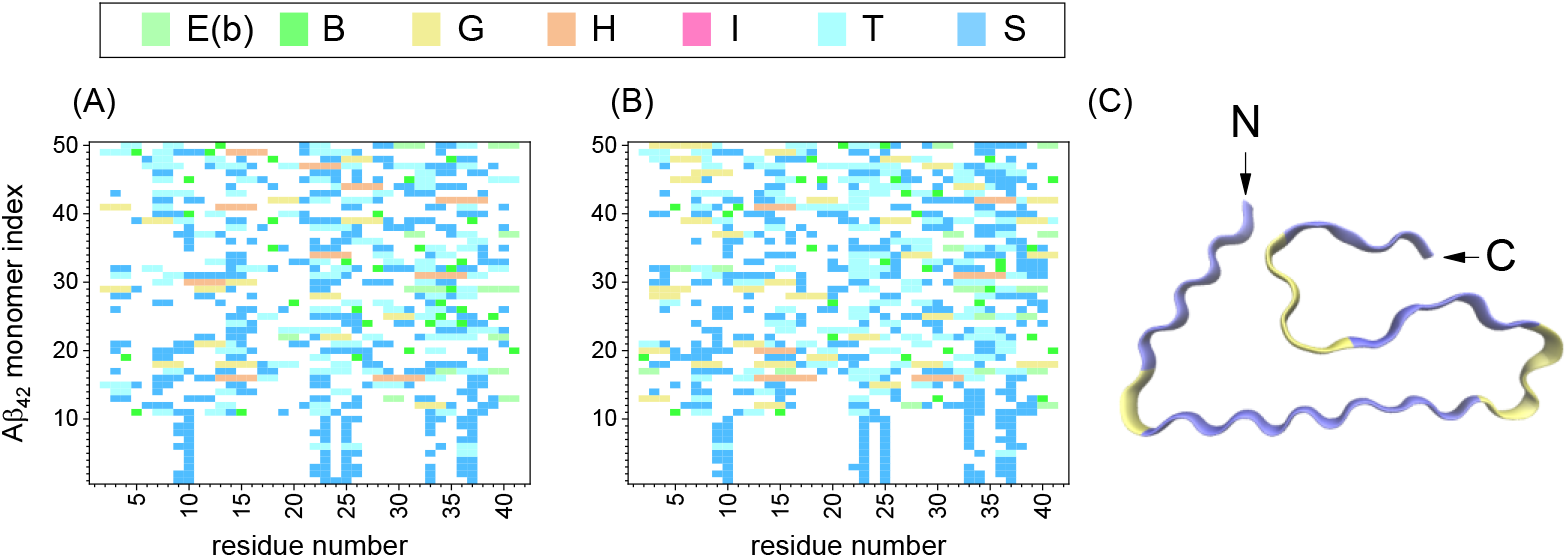
Secondary structure (SS) annotations for Aβ_42_ monomers. (A) Aβ_42_(M). (B) Aβ_42_(R|M). (C) Aβ_42_ monomer structure in fibril-growth competent form, the LS shape, which is found in the Aβ_42_ fibril structure registered with PDB entry: 5OQV. With regard to panels A and B, SS annotations are given as single letter format: E(b): extended β; B: isolated β; G: 3-10 helix; H: α helix; I: π(3-14) helix; T: turn; S: bend. In panel C, the letters ‘N’ and ‘C’ indicate N- and C-terminals of Aβ_42_ monomer, respectively. The following residues are involved in turn and bend formation and colored by yellow: Gly9, Thr10, Glu22, Asp23, Val24, Gly25, Ser26, Gly33, Leu34, Met35, Val36, Gly37.

Meanwhile, such SS formation features are not found in some of Aβ_42_ monomers forming the aggregate milieu. The turn and bent structures around 10^th^, 25^th^ and 37^th^ residues are not necessarily retained for each of the 40 Aβ_42_ monomers, indexed from 11 to 50. Instead, the remaining amino acid residues in each monomer shows additional turn and bent at the other regions, and also helix and strand formations in some cases.

The above analyses support the observation that these 40 Aβ_42_ monomers take forms deviated from the LS shape conformation^24^. We further confirmed this observation by examining the RMSd values to the reference structure, the Aβ_42_ monomer in LS shape conformation shown in **Figure 5C**. The Aβ_42_ monomers in Aβ_42_(5:5) have RMSd value of about 0.2 nm, while each of Aβ_42_ monomers forming Aβ_42_ aggregate milieus shows RMSd value greater than 1 nm (**Figure 6**). According to the above SS and RMSd analyses, we can say that the components of Aβ_42_ aggregate milieus, which are prepared by using the hybrid MC/MD simulations, are apparently structurally different from each of Aβ_42_ monomers in Aβ_42_(5:5). Thus, we can suppose that these Aβ_42_ monomers form off-pathway, fibril-growth incompetent Aβ_42_ aggregates rather than fibril-growth competent Aβ_42_ oligomer such as Aβ_42_(5:5).

**Figure 6.**
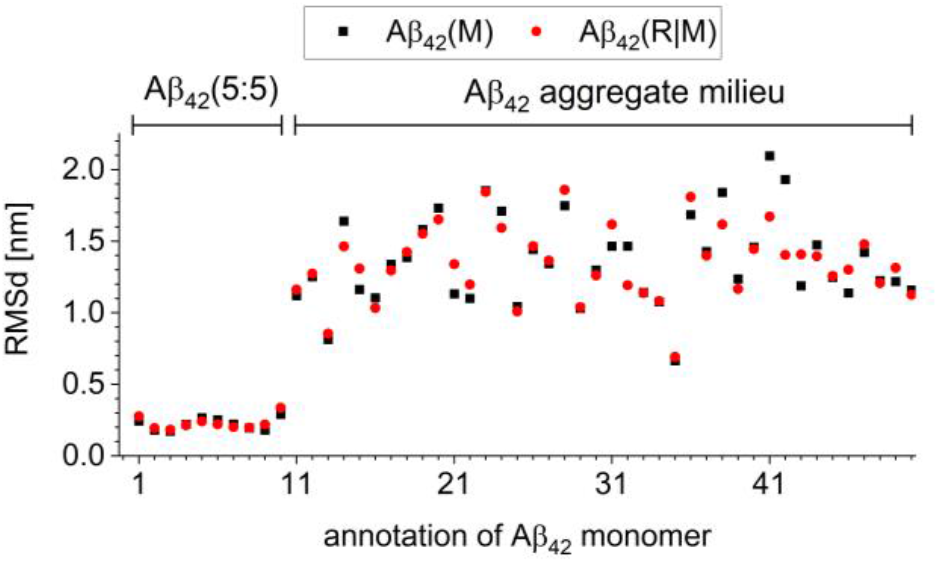
Root mean square deviation (RMSd) from the fibril-growth competent LS shape conformation (a monomer derived from the amyloid fibril deposited as PDB entry: 5OQV). Aβ_42_(M) and Aβ_42_(R|M) are shown by black and red symbols, respectively.

We investigated the number of Aβ_42_ monomer forming each Aβ_42_ aggregate milieu and classified them into two groups with regard to the presence and absence of atomic contacts with Aβ_42_(5:5). The results are summarized in **Table 3** and **Table 4** for Aβ_42_(M) and Aβ_42_(R|M), respectively. It is noted that an annotation with capital letter (*e*.*g*., A_1_) denotes an aggregate milieu which makes atomic contacts with any of monomers in Aβ_42_(5:5) and that with minuscule letter (*e*.*g*., a_1_) denotes the aggregate milieu without making atomic contacts with Aβ_42_(5:5).

**Table 3.** Classification of Aβ_42_ monomers (components 11-50) forming Aβ_42_ aggregate milieu in Aβ_42_(M) system. Aβ_42_ monomers in aggregate milieu, which make atomic contacts with any of monomers in Aβ_42_(5:5) (component 1-10), are highlighted by bold character.

| Aggregate | Component |
| --- | --- |
| A<sub>1</sub> | 14,15,16,17,21,22,23,24,25,26,27,40,**42**,43,**44**,45,46,47 |
| A<sub>2</sub> | **13**,20,**28,29**,32,33,34,35,36,37,38,**48** |
| a<sub>1</sub> | 12,30,31,49 |
| A<sub>3</sub> | 18,**19** |
| a<sub>2</sub> | 11,50 |
| a<sub>3</sub> | 39 |
| a<sub>4</sub> | 41 |

**Table 4.** Classification of Aβ_42_ monomers (components 11-50) forming Aβ_42_ aggregate milieu in Aβ_42_(R|M) system. Aβ_42_ monomers in aggregate milieu, which make atomic contacts with any of monomers in Aβ_42_(5:5) (component 1-10), are highlighted by bold character. The subscript ‘r’ denotes relaxation by the 100-ns NPT MD simulation.

| Aggregate | Component |
| --- | --- |
| A<sub>1r</sub> | 14,15,16,17,18,**19**,24,25,26,27,30,31,40,**41,42**,43,**44**,45,46,49 |
| a<sub>1r</sub> | 12,35,36,37,38,39 |
| A<sub>2r</sub> | 20,**28**,29,32,33,34 |
| a<sub>2r</sub> | 11,21,22,23,50 |
| A<sub>3r</sub> | **13**,48 |

**Table 5.** Buried solvent accessible surface area for Aβ_42_ monomers in the fibril growth edge of Aβ_42_(5:5) [unit: nm^2^]. Monomers in Aβ_42_(5:5) are annotated as in the case of **Figure 1**

**Figure 1.**
| monomer in Aβ<sub>42</sub>(5:5) | Aβ<sub>42</sub>(M) | Aβ<sub>42</sub>(R\|M) |
| --- | --- | --- |
| S1 | 7.22 | 6.88 |
| S5 | 7.73 | 6.56 |
| S6 | 0 | 0 |
| S10 | 9.01 | 17.07 |

For both of Aβ_42_(M) and Aβ_42_(R|M), Aβ_42_(5:5) makes contacts with seven or six Aβ_42_ monomers in Aβ_42_ aggregate milieu (annotated by bold characters in **Tables 3** and **4**), respectively. This observation indicates that Aβ_42_(5:5) is not encompassed by Aβ_42_ aggregate milieu, but makes local contacts with Aβ_42_ aggregates. In other words, Aβ_42_(5:5) forms a pairwise complex with an Aβ_42_ aggregate.

We also evaluated the global shape of a component of Aβ_42_ aggregate milieu, which has the largest number of Aβ_42_ monomer, A_1_ and A_1r_ for Aβ_42_(M) and Aβ_42_(R|M), respectively. The values of hydrodynamic radius (*R*_*hyd*_, which is defined in **eqn. 1**) are 12 and 25 nm for Aβ_42_(M) and Aβ_42_(R|M), respectively. According to the earlier experimental studies, a micelle-like Aβ_42_ aggregate consists of 25 to 50 Aβ_42_ monomers^18,19^ and shows value of *R*_*hyd*_ in range from 2.4 to 11.4 nm^18^. Thus, we could suppose our simulation-derived aggregate species are not such micelle-like aggregates. These aggregates are less condensed spatially than a micelle-like Aβ_42_ aggregate but are rather extended in the space, as indicated by analyses of *R*_*hyd*_. It is thus noted that we here discuss new facet of Aβ_42_ aggregate milieu effect on stable formation of Aβ_42_ growth nuclei below.

Among the four Aβ_42_ monomers at the fibril growth edge (S1, S5, S6 and S10) of Aβ_42_(5:5), the monomers except for S6 make contacts with any of the Aβ_42_ aggregates (see **Figure 3**). For the three monomers, we analyzed solvent accessible surface area (SASA) covered by an amyloid aggregate and found that S10 has the greatest buried SASA value of the three Aβ_42_ monomers for both Aβ_42_(M) and Aβ_42_(R|M). Magnitude of the buried SASA possibly correlates with energetic stability of intermolecular interaction. We then supposed that S10 relatively strongly interacts with Aβ_42_ aggregates compared with S1 and S5 and considered the S10 for the free energy calculations of monomer dissociation from the fibril growth edge.

### Aβ_42_ aggregate milieu suppresses Aβ_42_ monomer dissociation from the fibril growth edge by anchoring the monomer to Aβ_42_ growth nuclei

We will examine the effects of Aβ_42_ aggregate milieu on Aβ_42_ monomer dissociation processes as follows. First, we performed independent 50 SMD simulations for the Aβ_42_ monomer (S10) dissociation from the fibril growth edge of the Aβ_42_(5:5). Each of the three snapshot structures, Aβ_42_(M), Aβ_42_(P) and Aβ_42_(R|M), was employed as the set of the initial atomic coordinates. It is noted that the simulated results are annotated by the corresponding Aβ_42_ snapshot structure, namely, Aβ_42_(M), Aβ_42_(P) or Aβ_42_(R|M), hereafter.

**Figure 7** shows the trajectory of center of mass of the dissociating Aβ_42_ monomer, S10. A set of SMD simulations starting from Aβ_42_(P) shows a broader distribution of the center of mass (COM) of the dissociating Aβ_42_ monomer than those from Aβ_42_(M) and Aβ_42_(R|M). We can obtain this observation by comparing **Fig. 7A-C** with **Fig. 7D-F** and **Fig. 7G-I**.

**Figure 7.**
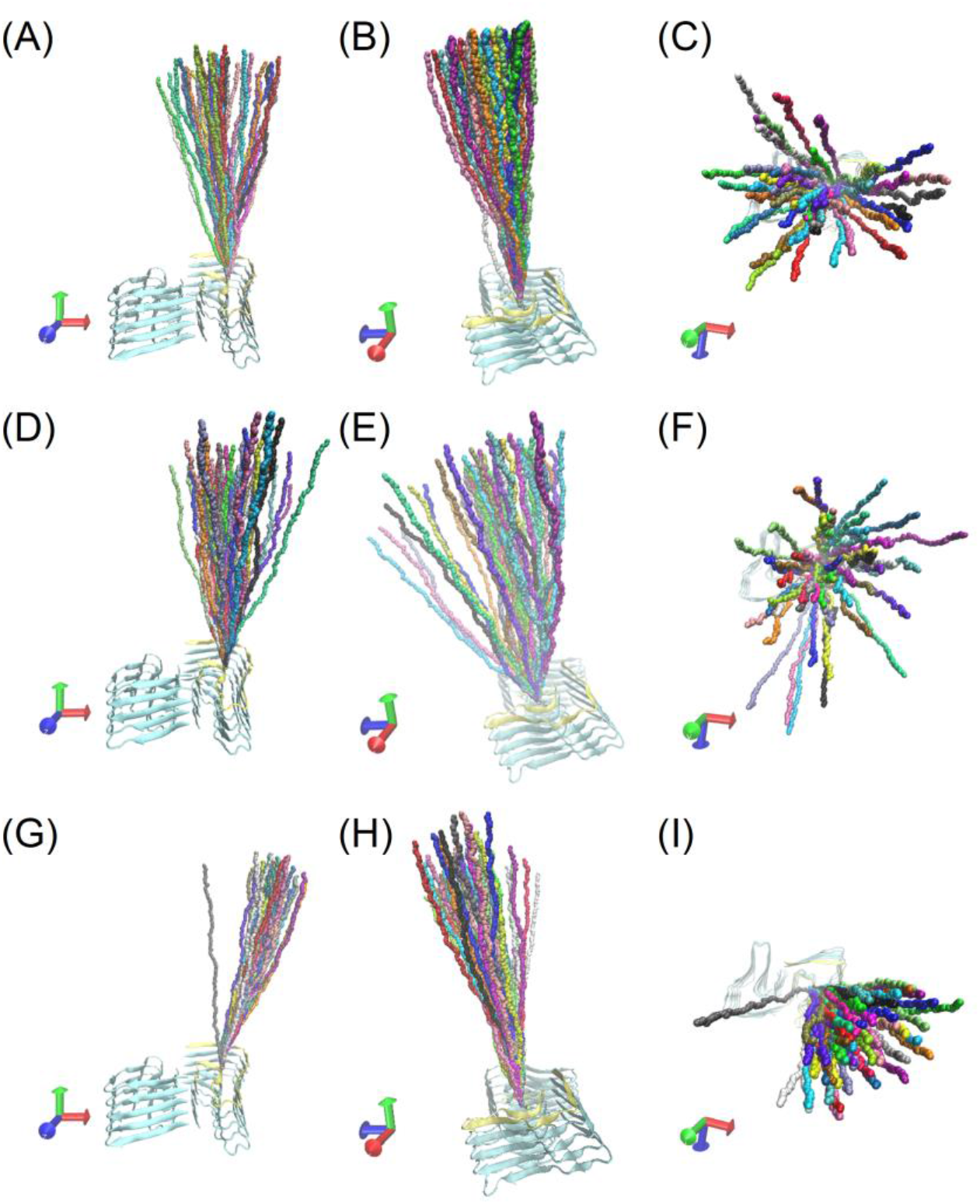
Dissociating pathways of Aβ_42_ monomer from the growth edge. (A)-(C) Aβ_42_(M). (D)-(F) Aβ_42_(P). (G)-(I) Aβ_42_(R|M). Center of mass (COM) of the dissociating Aβ_42_ monomer is represented by colored sphere. Each of the 50 COM trajectories is distinguished by the different color. The dissociation-attempted Aβ_42_ monomer in Aβ_42_(5:5), S10, and the remaining of Aβ_42_(5:5) are represented by yellow and blue ribbon, respectively.

We confirm the above observation by quantitatively characterizing the spatial expansion of the COM distributions with an effective radius at each time point, which is defined in **eqn. 3**. As shown in **Fig. 8**, along the time course of the SMD simulation, the effective radius for Aβ_42_(M) and Aβ_42_(R|M) is smaller than that for Aβ_42_(P). It can thus be said that the presence of amyloid aggregate milieu spatially restricts dissociation pathways of Aβ_42_ monomer from the edge of growth axis.

**Figure 8.**
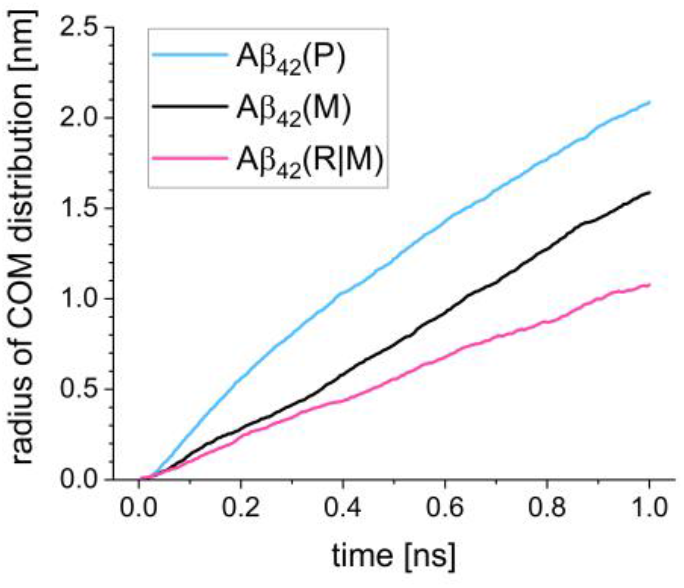
Spatial expansion of Aβ_42_ monomer dissociating trajectory, estimated by the effective radius of dissociating trajectory set (see **eqn. 3** for the definition).

The narrowed distributions of Aβ_42_ monomer dissociation pathways remind us of entropic contribution of Aβ_42_ aggregate milieu to stable formation of Aβ_42_ growth nuclei. The Aβ_42_ aggregate milieu has much larger inertia than water molecules which can move flexibly in the system, so that the milieu could more stably reside at the same place than water molecules and could reduce the space which the dissociation monomer can access. Therefore, it is straightforward expectation that such a reduction of accessible space prevents Aβ_42_ monomer dissociation with increase of entropic costs.

We tested such entropic effects of Aβ_42_ aggregate milieu on energetic stability of Aβ_42_ monomer binding to the fibril growth axis by calculating free energy profiles of Aβ_42_ monomer dissociation from the fibril growth edge of Aβ_42_(5:5). For each of the three systems, we randomly selected 6 SMD simulations among the 50 ones, and used the six SMD trajectories for individual USMD simulations. As remarked in Materials and Method section, 6 is the minimum to make 200 resampling for bootstrap error estimation; the resampling number of 200 is the commonly used number in this field for bootstrap error estimation^36^.

Interestingly, there are no significant difference between Aβ_42_(M) and Aβ_42_(P) in the activation barrier of Aβ_42_ monomer dissociation (see black lines in **Fig. 9A** and **9B**, respectively). The activation barrier of Aβ_42_(P) is 35.2 ± 5.2 kJ/mol, while that of Aβ_42_(M) is 39.6 ± 9.6 kJ/mol. As discussed with regard to **Fig. 8**, the presence of Aβ_42_ aggregate milieu restricted accessible space for the Aβ_42_ monomer, S10, upon the dissociation from the growth edge, then suggesting an entropic cost to increase free energy barrier of the Aβ_42_ monomer dissociation process. However, our free energy calculations result that the entropic effect discussed in **Fig. 8** does not significantly contributes to Aβ_42_ monomer binding to the growth edge of Aβ_42_ oligomer. We could say that an Aβ_42_ aggregate just surrounding an Aβ_42_ growth nuclei does not necessarily work for stable formation of Aβ_42_ growth nuclei.

**Figure 9.**
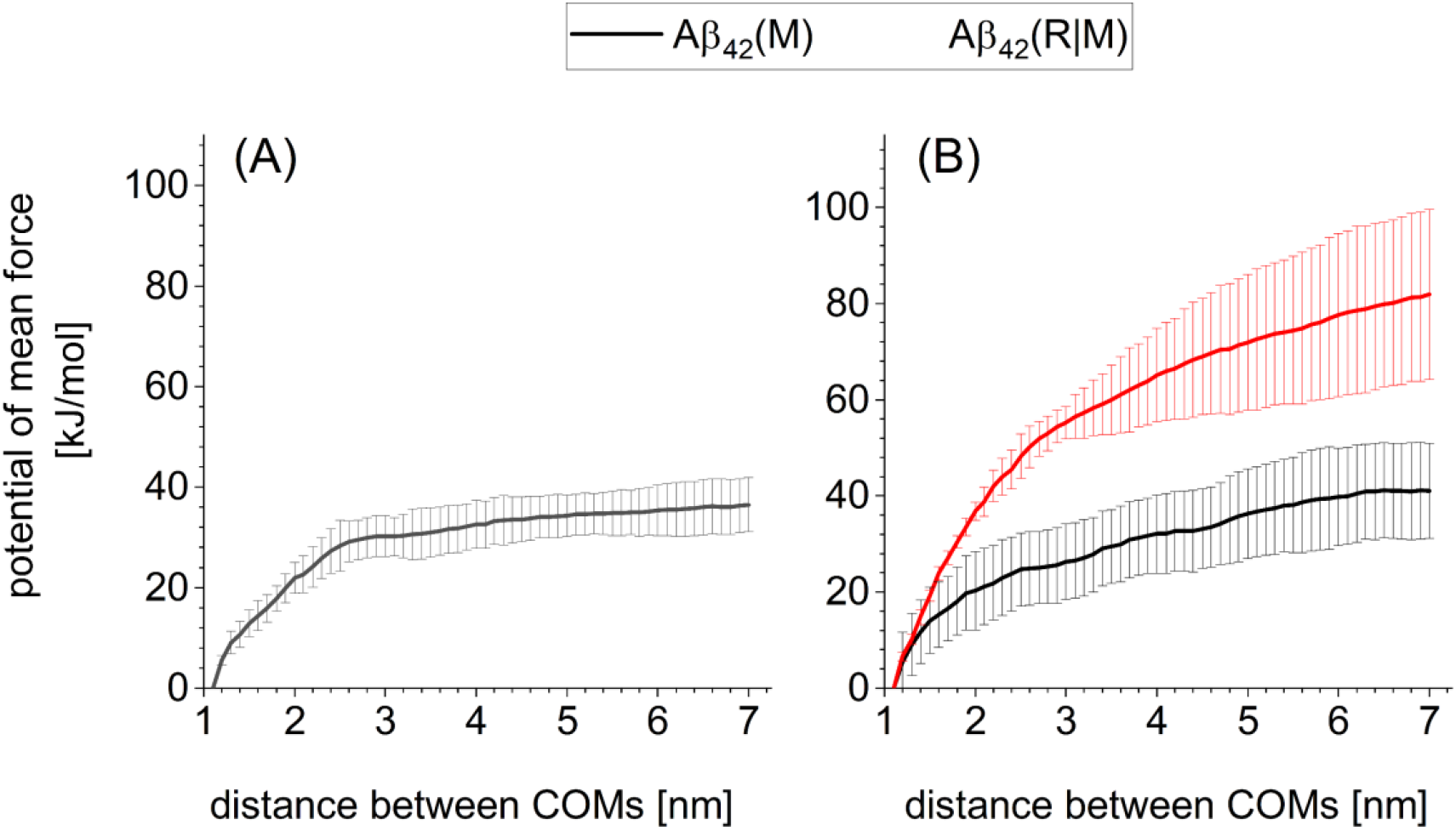
Potential of mean force for Aβ_42_ monomer dissociation from the fibril growth edge of pentameric oligomer. (A) Aβ_42_(5:5) in aqueous solution, Aβ_42_(P). (B) Aβ_42_(5:5) in Aβ_42_ crowding milieu. In the panel B, black and red lines are for Aβ_42_(M) and Aβ_42_(R|M), respectively.

Meanwhile, there are significant difference between Aβ_42_(R|M) and the other two Aβ_42_ systems in Aβ_42_ monomer binding to the growth edge: the activation barrier is 82.4 ± 17.2 kcal/mol (see red line in **Fig. 9B**). The significant difference between Aβ_42_(M) and Aβ_42_(R|M) suggests the microscopic origin of roles of Aβ_42_ amyloid aggregate in thermodynamically stabilizing Aβ_42_ monomer binding, that is, enthalpic stabilization with direct atomistic interaction with an Aβ_42_ amyloid growth nuclei such as Aβ_42_(5:5). It is worthwhile noting that in Aβ_42_(R|M), a monomer in the amyloid aggregate milieu, S19, makes atomic contacts with both the dissociating monomer, S10, and the remaining of Aβ_42_(5:5) (**Fig. 10**). This observation lets us suppose that the S19 anchors S10 on the remaining of Aβ_42_(5:5) and cooperatively suppresses S10 dissociation from the growth edge.

**Figure 10.**
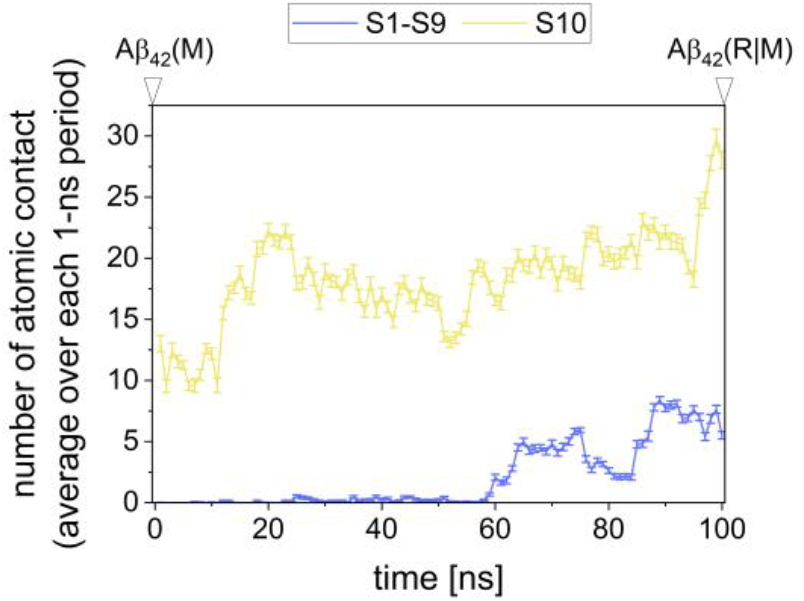
Atomic contacts of S19 with S10 and the remaining of Aβ_42_(5:5), nine monomers from S1 to S9, shown by yellow and blue lines, respectively. The atomic contact number is averaged over each 1-ns period and the error bars indicate 95% confidence interval.

If such anchoring effect works for suppression of S10 dissociation, S19 in Aβ_42_(R|M) maybe more strongly interacts with Aβ_42_(5:5) than in Aβ_42_(M). To verify this speculation, we examined the dissociation free energy landscape for S19. As the initial structures for free energy landscape calculations, we used the hybrid MC/MD-derived configuration and the subsequent 100-ns NPT MD derived configuration, which are discussed in **Figure 4A** and **4B** as Aβ_42_(M) and Aβ_42_(R|M), respectively.

By comparing the free energy landscapes between Aβ_42_(R|M) and Aβ_42_(M), we can find that Aβ_42_(R|M) has a greater PMF value than Aβ_42_(M) (see **Fig. 11**). As for Aβ_42_(M), the height of activation barrier is less than 8 kJ/mol. Meanwhile, Aβ_42_(R|M) shows an up-hill shape of the free energy landscape and has ca 32 kJ/mol at 8 nm. The S19 more stably interacts with the Aβ_42_(5:5) in Aβ_42_(R|M), due to the greater number of atomic contacts (see **Fig. 10**), than in Aβ_42_(M). S19 of Aβ_42_(R|M) appears to loosely bind to Aβ_42_(5:5) with relatively weak interaction in comparison with the S10 interaction with Aβ_42_(5:5), while such loose interaction with Aβ_42_(5:5) may be sufficient to cooperatively suppress Aβ_42_ monomer dissociation from the growth edge (see **Fig. 9B**).

**Figure 11.**
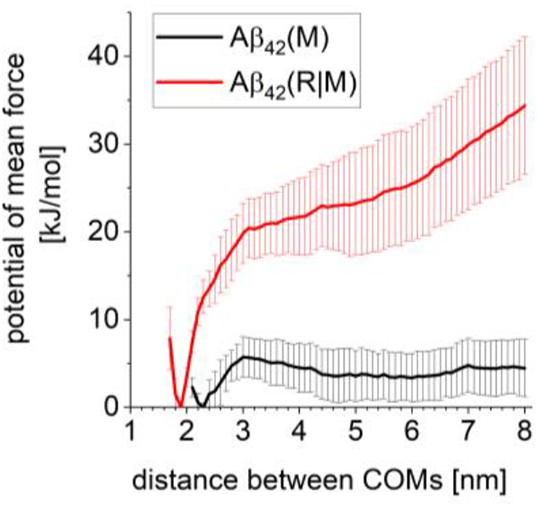
Potential of mean force for aggregate milieu component Aβ_42_ monomer (S19) dissociation from the Aβ_42_(5:5). Black and red lines are for Aβ_42_(M) and Aβ_42_(R|M), respectively.

## Concluding Remarks

In the present study, we discussed roles of the Aβ_42_ aggregate milieu in stable formation of Aβ_42_ growth nuclei by employing atomistic molecular dynamics simulations. We found that off-pathway, fibril-growth incompetent oligomeric Aβ_42_ aggregates directly interact with both dissociating Aβ_42_ monomer and the remaining of the growth nuclei and suppress the monomer dissociation by significant increase of the height of activation barrier in the process. We can suppose that such Aβ_42_ aggregates enthalpically stabilizes and progresses growth nuclei formation by loosely but cooperatively binding to the growth nuclei and dissociation-prone monomer at the edge of the growth nuclei. Although our observation for the effect of Aβ_42_ aggregate crowding milieu is obtained from one hybrid MC/MD simulation-derived Aβ_42_ aggregates structure, the microscopic mechanism proposed on the basis of the observation appears to be independent of specific atomistic details of aggregate crowding milieus. Thus, we may suppose that it is worthwhile to consider the mechanism for other amyloid aggregate milieus.

It is also worthwhile to remark that our simulation-derived Aβ_42_ aggregates do not take a micelle-like form but rather appear to take unstructured and spatially expanded configurations. It has been considered that micelle-like Aβ_42_ aggregates play important roles in Aβ_42_ fibril formation^17-20^. Besides this conventional observation, we newly propose that off-pathway, fibril-growth incompetent, non-micelle-like Aβ_42_ aggregates can contribute to stable formation of Aβ_42_ growth nuclei by suppressing Aβ_42_ monomer dissociation. It is reported that ATP molecules in physiological concentration nonspecifically dissolve protein aggregates including Aβ_42_ aggregates^26,38^. Recalling the roles of Aβ_42_ aggregate milieu in stable formation of Aβ_42_ aggregate growth nuclei, we speculate that ATP molecules suppress formations of both fibril-like aggregates and aggregate milieus simultaneously to preserve healthy cell conditions. Nonetheless, it is beyond the scope of this study to test the speculation, thus being left for future study.

## Abbreviations

(Aβ_42_): amyloid protein, amyloid-β (1-42)
(cryo-EM): cryogenic electron microscopy
(MM): molecular mechanics
(MD): molecular dynamics
(SMD): steered molecular dynamics
(USMD): umbrella sampling molecular dynamics
(PMF): potential of mean force
(HB): hydrogen bond
(SASA): solvent accessible surface area

## Supporting Information

The Supporting Information is available. Detailed procedures for unbiased MD, SMD and USMD simulations, Figures and Tables for analyses of these simulations.

## Acknowledgements

This work was supported by a Grant-Aid for Scientific Research on Innovative Areas “Chemistry for Multimolecular Crowding Biosystems” (JSPS KAKENHI Grand No. JP17H06353) and by MEXT Quantum Leap Flagship Program (Grant No. JPMXS0120330644).

