## Supporting Information for "Non-micelle-like Amyloid Aggregate Stabilizes Amyloid β (1-42) Growth Nuclei Formation"

^*^Ikuo Kurisaki

^*^Shigenori Tanaka

**SI-1 Construction of amyloid-β (1-42) 25 mer dimer**

We used the cryo-electron microscopy (cryo-EM) structure (PDB entry: 5OQV^1^) to construct Aβ_42_ 25 mer protomer dimer, Aβ_42_(25:25). Aβ_42_(4:4) derived from the cryo-EM structure was used as the initial structure. Larger Aβ_42_(N:N) were generated by superposition of two Aβ_42_(4:4) on the edges of the protomer dimers and following deletion of overlapped monomer pairs. This procedure was similar to that discussed in our previous study^2^ and was repeated to obtain the Aβ_42_ 25 mer dimer, which was performed by using root mean square fit with cpptraj module in AmberTools^3^.

**SI-2 Unbiased MD simulation for Aβ_42_(25:25) system**

For the Aβ_42_(25:25) system, the atomic coordinates of water and K^+^ molecules were energetically relaxed by the following molecular mechanics (MM) and molecular dynamics (MD) simulations. In each of the following MM and MD simulations, the atomic coordinates of non-hydrogen atoms in the Aβ_42_(25:25) were restrained by the harmonic potential with force constant of 0.4184 kJ/mol/nm^2^ around the initial atomic coordinates.

First, steric clashes in the system were removed by MM simulation, which consists of 1000 steps of the steepest descent method followed by 49000 steps of the conjugate gradient method. Then the system temperature and density were relaxed through the following five MD simulations: NVT (0.001 to 1 K, 0.1 ps) → NVT (1 K, 0.1 ps) → NVT (1 to 300 K, 20 ps) → NVT (300 K, 20 ps) → NPT (300 K, 300 ps, 1 bar).

The first two NVT MD simulations and the other MD simulations were performed using 0.01 fs and 2 fs for the time step of integration, respectively. The first and second NVT MD simulations were performed using Berendsen thermostat^4^ with a 0.001 ps of coupling constant. Meanwhile the following three simulations were performed using Langevin thermostat with 1-ps^-1^ of collision coefficient. In the first NVT MD simulation, the reference temperature was linearly increased along the time-course. In the NPT MD simulation, the system pressure was regulated with Monte Carlo barostat, where the system volume change was attempted by every 100 steps. Each set of initial atomic velocities was randomly assigned from the Maxwellian distribution at 0.001 K.

Using the density-relaxed atomic coordinates derived from the above simulation, an Aβ_42_(25:25) conformation also was structurally relaxed in aqueous solution through the following 7-step MD simulations: NVT (0.001 to 1 K, 0.1 ps, 0.4184 kJ/mol/nm^2^) → NVT (1 to 300 K, 0.1 ps, 0.4184 kJ/mol/nm^2^) → NVT (300 K, 10 ps, 0.4184 kJ/mol/nm^2^) → NVT (300 K, 40 ps, 0.2092 kJ/mol/nm^2^) → NVT (300 K, 40 ps, 0.04184 kJ/mol/nm^2^) → NVT (300 K, 40 ps) → NPT (300 K, 1 bar, 10 ns). The first two NVT MD simulations and the other MD simulations were performed using 0.01 fs and 2 fs for the time step of integration, respectively. In the first two NVT simulations, the reference temperature was linearly increased along the time-course. In the first 5 steps, non-hydrogen atoms in Aβ_42_ protomer dimer were positionally restrained by the harmonic potential around the initial atomic coordinates. In each NVT simulation, temperature was regulated using Langevin thermostat with 1-ps^-1^ collision coefficient. In the last 10-ns NPT simulation, temperature and pressure were regulated by Berendsen thermostat^4^ with a 5-ps coupling constant, and Monte Carlo barostat, where system volume change was attempted by every 100 steps, respectively. The initial atomic velocities were randomly assigned from the Maxwellian distribution at 0.001 K. The snapshot structure obtained from the 10-ns NPT MD simulation procedure was employed for the following SMD simulations.

**S3. Hybrid (configuration bias) MC/MD simulation scheme**

The simulation scheme for consecutive Aβ_42_ monomer dissociation is similar to that disassembly of serum amyloid P component protein 5 mer, which was considered in our previous study^5^. We here employed unbiased and SMD simulations to generate *old* and *new* sets of atomic configurations of molecular systems, respectively. Inter-Aβ_42_ monomer dissociation reactions are accelerated with SMD simulations. Detailed simulation conditions of MD and SMD-based configuration generation are given in the next subsection.

1. Generate *M* trial configurations
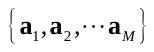
 by an unbiased MD simulation and calculate bias energy *u^bias^*, which will be specified later, for each configuration.
2. Assign the last snapshot obtained from the simulation (
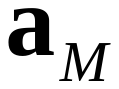
) as an old configuration, denoted by
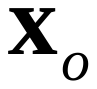
 and define the Rosenbluth factor^6^

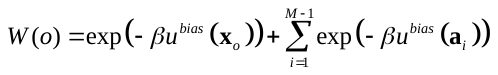
 (1)

*β* denotes inverse of k_B_T, where k_B_ and T are Boltzmann constant and system temperature, respectively. In the following step 3, new configurations are generated by starting from
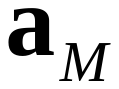
, so that we assigned this configuration to
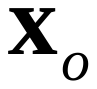
 by recalling conventional cbMC schemes^6^.

3. Generate *M* trial configurations
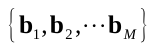
 by a steered MD simulation starting from the old configuration
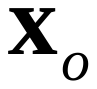
 and calculate bias energy *u^bias^* for each configuration.
4. Define the Rosenbluth factor

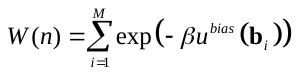
 (2)

and select one configuration among
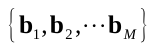
, denoted by
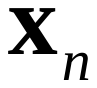
, with a probability

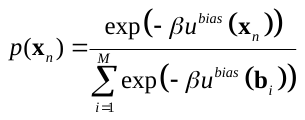
 (3)

5. The configuration change is accepted with a probability

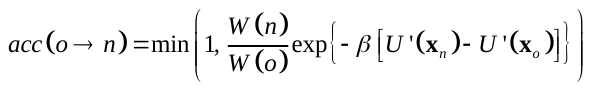
 (4)

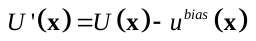
 (5)

where
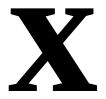
 and
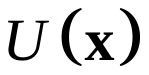
 denote a configuration and the potential energy function of the system with configuration of
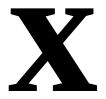
, respectively.

Considering that configurations of the system are generated by moving all atoms simultaneously with molecular dynamics method, we used the Rosenbluth factor in the forms given in Eq. 1 and Eq. 2.

A multimeric protein complex disassembly process should be accompanied by gradual breakage of inter-subunit native contacts (NC). Then we defined the bias energy as a function of the total number of inter-monomer NC formed among Aβ_42_ monomers (*n_NC_*), where the initial atomic coordinates are used as the reference structure:

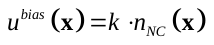
 (6)

The coefficient *k* is set to *β*^−1^ at 300 K, that is 0.6 [kcal/mol] as in the case of our previous study^5^. A set of native contacts at Aβ_42_ monomer binding interface was defined by using the initial atomic coordinates of Aβ_42_(25:25) for hybrid MC/MD simulations. All hybrid MC/MD simulations were performed with employing an in-house MC/MD simulation interface during MD simulations with Amber17^3^.

**S4. Configuration generation by using unbiased MD and steered MD simulations**

Configurations of the Aβ_42_ system, denoted by
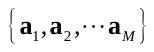
, were sampled by using an unbiased NPT MD simulation (300 K, 1 bar). Meanwhile, configurations of the Aβ_42_ system undergoing a partial dissociation, denoted by
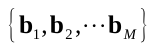
, were sampled by using a steered molecular dynamics (SMD) simulation under the NPT condition (300 K, 1 bar), starting from the snapshot structure obtained from the MD simulation. The simulation length is 100 ps for each of MD and SMD simulations.

In every hybrid MC/MD cycle at each dissociation stage, we randomly chose a pair of monomers from the candidate list (see **Table 1**), and employed the distance between centers of mass of the chosen Aβ_42_ monomer pair as the reaction coordinate of SMD simulation. A center of gravity for an Aβ_42_ monomer is calculated by using all C_α_ atoms in the monomer. In an SMD simulation, a target value of the distance was set to *d*_0_ + Δ*d*, where *d*_0_ and Δ*d* are an initial value of the distance and a random integer in the range of 8 to 12, respectively. The harmonic potential with the force constant of 4.184 J/mol/nm^2^ was imposed on the reaction coordinate.

The system temperature and pressure were regulated by Langevin thermostat with a 1-ps^−1^ collision frequency, and Monte Carlo barostat with attempt of system volume change by every 100 steps, respectively. The trajectory was recorded every 10-ps interval. In each of MD and SMD simulations, a set of initial atomic velocities was calculated from products of integration time step and atomic forces acting on the atoms.

**SI-5 Calculation of potential of mean force**

Steered molecular dynamics (SMD) simulations were performed to prepare the initial atomic coordinates for each window of umbrella sampling (US) MD simulations by considering the reaction coordinate defined in the main text. The value of reaction coordinate was gradually changed through the SMD simulations by imposing the harmonic potential with force constant of 4.184 kJ/mol/nm^2^. The target distance of SMD simulations was set to 10 nm. The SMD simulation was executed for 1 ns under NPT condition (300 K; 1 bar). The trajectory was recorded every 5-ps interval. Temperature and pressure were regulated using Langevin thermostat with a 1-ps^−1^ of collision coefficient and Berendsen barostat^4^ with a 5-ps coupling constant, respectively. A set of initial atomic velocities were taken over from the previous MD simulation. Using each SMD trajectory, we prepared snapshot structures for a given set of umbrella windows (Tables **S1**, **S2** and **S3** for Aβ(M), Aβ(P) and Aβ(R|M) in S10 dissociation, respectively; Tables **S4** and **S5** for Aβ(M) and Aβ(R|M) in S19 dissociation, respectively).

For each of the three systems, the S10 dissociation procedure was repeated 50 times and randomly selected 6 simulations to perform USMD simulations. This is due to reduction of computational cost to consider huge molecular systems: the sample size of 6 is minimum requisite to give bootstrap-error estimation of 200 trials, which is used in the earlier studies^7^. As for S19 dissociation, we simply performed 6 independent SMD simulations for each of the two systems, because we do not have to examine dissociation pathways.

Using each initial atomic coordinates derived from the SMD simulations, we performed the relaxation simulation and the following USMD simulation. The relaxation simulation consists of 5 steps: NVT (0.001 to 1.0 K, 0.1 ps, 4.184 kJ/mol/nm^2^) → NVT (1.0 to 300 K, 0.1 ps, 4.184 kJ/mol/nm^2^) → NVT (300 K, 40 ps, 0.4184 kJ/mol/nm^2^) → NVT (300 K, 40 ps, 0.2092 kJ/mol/nm^2^) → NVT (300 K, 40 ps, 0.04184 kJ/mol/nm^2^). The initial atomic velocities were randomly assigned from the Maxwellian distribution at 0.001 K in the first step. Backbone heavy atoms (Cα, C, N, O) of Aβ_42_ protomer dimer were restrained by the harmonic potential around the initial atomic coordinates. In each of the first two NVT-MD simulations, the reference temperature was linearly increased along the time-course. Then, several nano-second USMD simulation was executed under NVT condition (300 K). For each system, the USMD simulation time length is determined by evaluating the convergence of PMF (*see* Panels A-C in **Fig. S1**). The reaction coordinate of these USMD simulations is the same as that for the SMD simulations. In an USMD simulation, temperature was regulated using Langevin thermostat with 1-ps^-1^ collision coefficient, and the interatomic distance for reaction coordinate was recorded every 1-ps interval.

Using 6 sets of USMD simulations, we made a complete histogram spanning the reaction coordinate from 1 nm to 8 nm. For each set of USMD Simulations, we confirmed 2.5 % or greater overlap in sampling of neighboring USMD windows. Then the complete histogram was employed to compute a potential of mean force, where width of bin was set to 0.1 nm.

The S19 dissociation simulations were carried out as in the case of S10 dissociation, while the reaction coordinate is selected for the simulations (see the main text). For each system, the USMD simulation time length is determined by evaluating the convergence of PMF (*see* Panels D and E in **Fig. S1**)

**Supporting Tables**

**Table S1.** Equilibrium positions of biased potentials and corresponding force constants for S10 monomer dissociation umbrella sampling molecular dynamics simulations, for Aβ_42_(M) system.

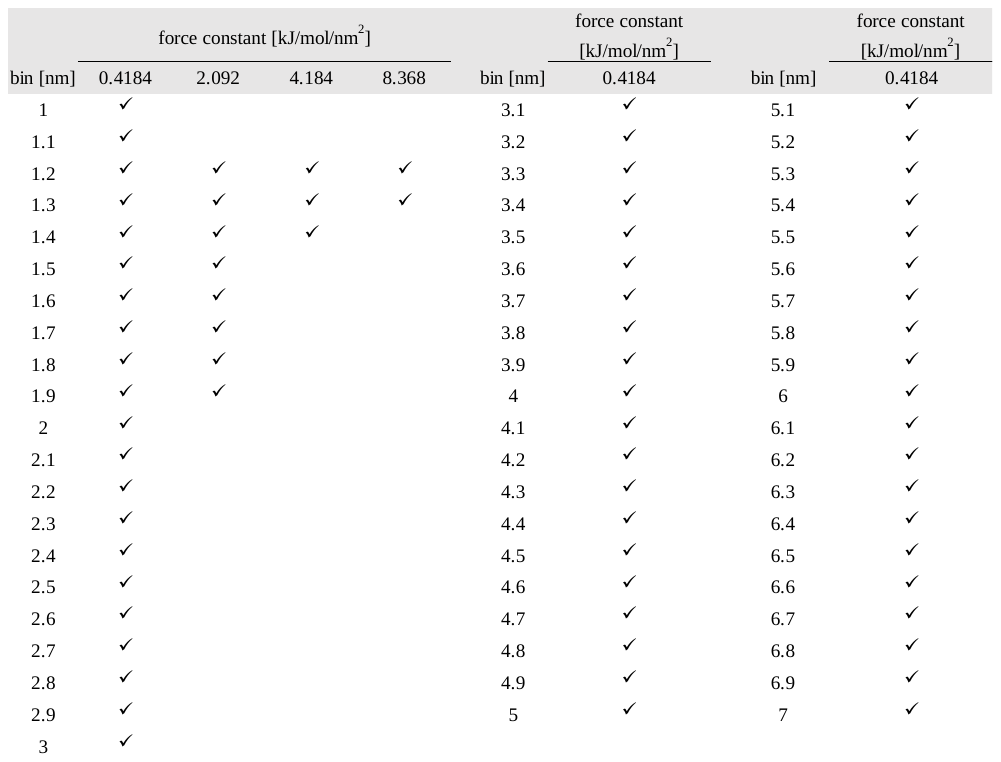

**Table S2.** Equilibrium positions of biased potentials and corresponding force constants for S10 monomer dissociation umbrella sampling molecular dynamics simulations, for Aβ_42_(P) system.

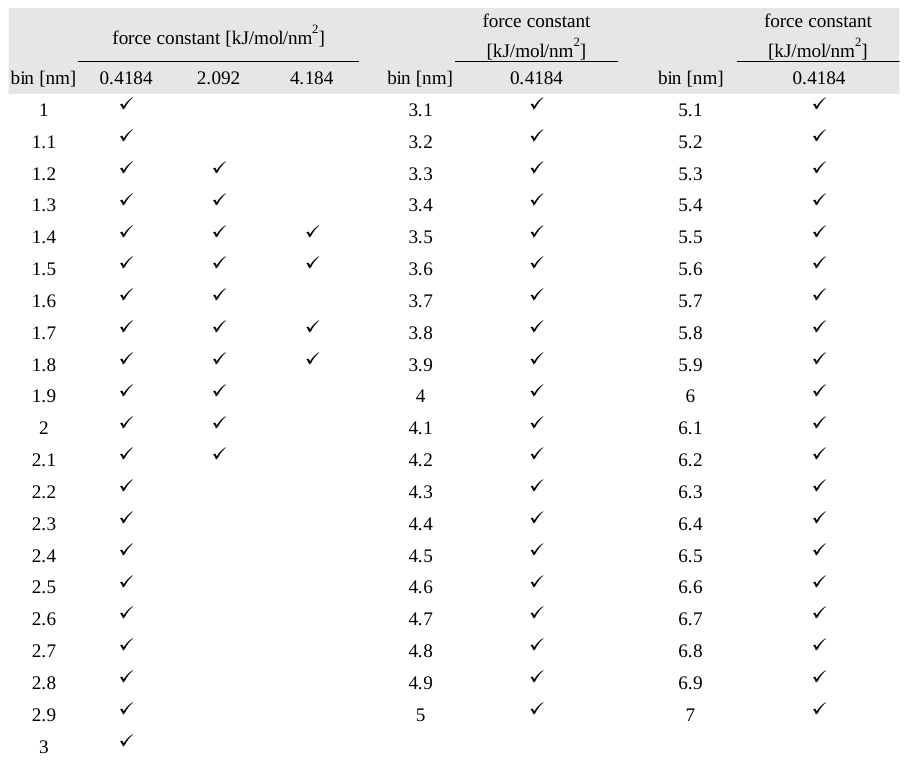

**Table S3.** Equilibrium positions of biased potentials and corresponding force constants for S10 monomer dissociation umbrella sampling molecular dynamics simulations, for Aβ_42_(M|R) system.

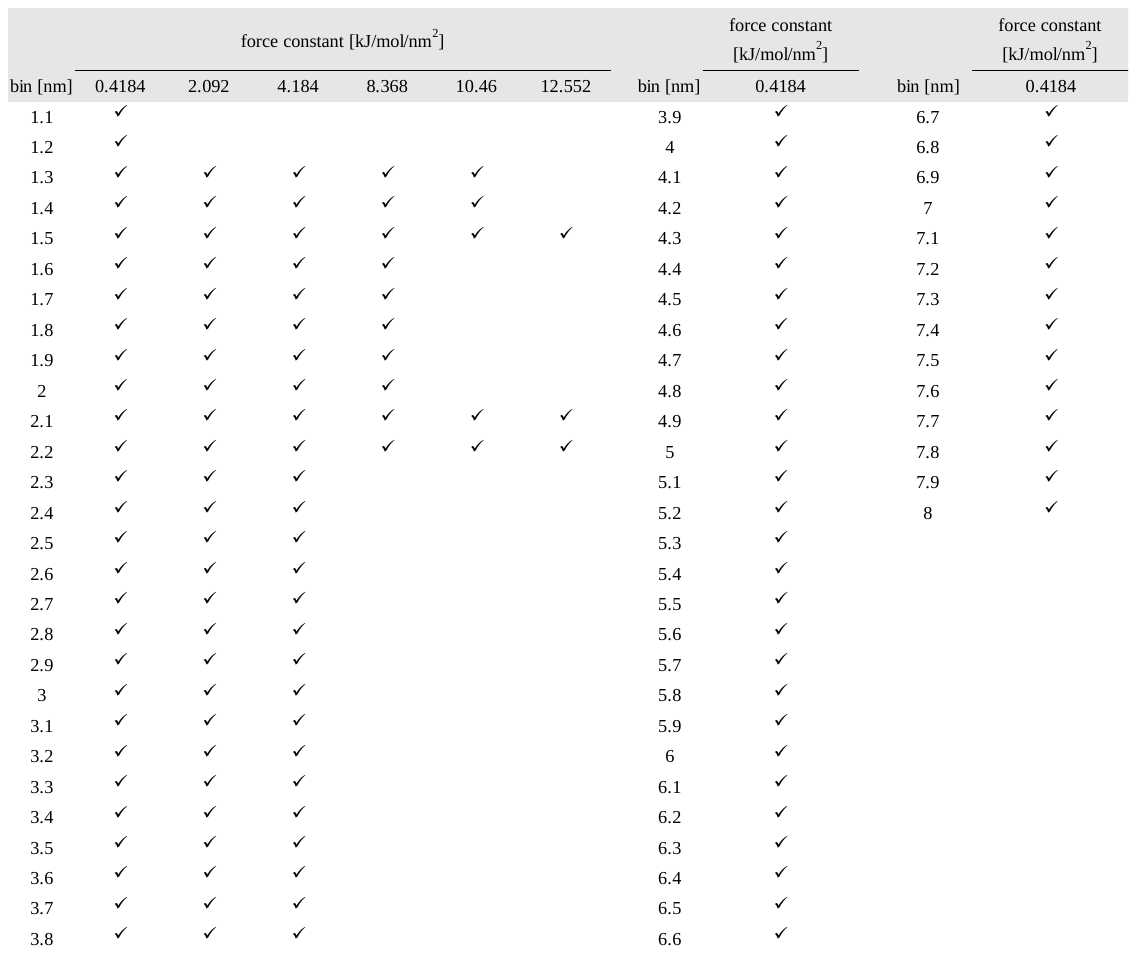

**Table S4.** Equilibrium positions of biased potentials and corresponding force constants for S19 monomer dissociation umbrella sampling molecular dynamics simulations for Aβ_42_(M) system.

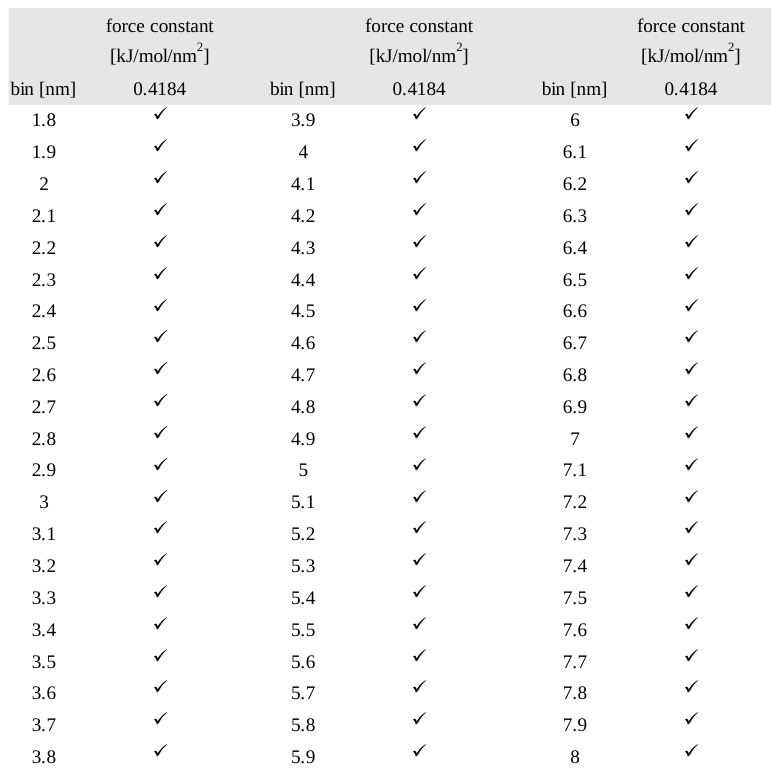

**Table S4.** Equilibrium positions of biased potentials and corresponding force constants for S19 monomer dissociation umbrella sampling molecular dynamics simulations for Aβ_42_(R|M) system.

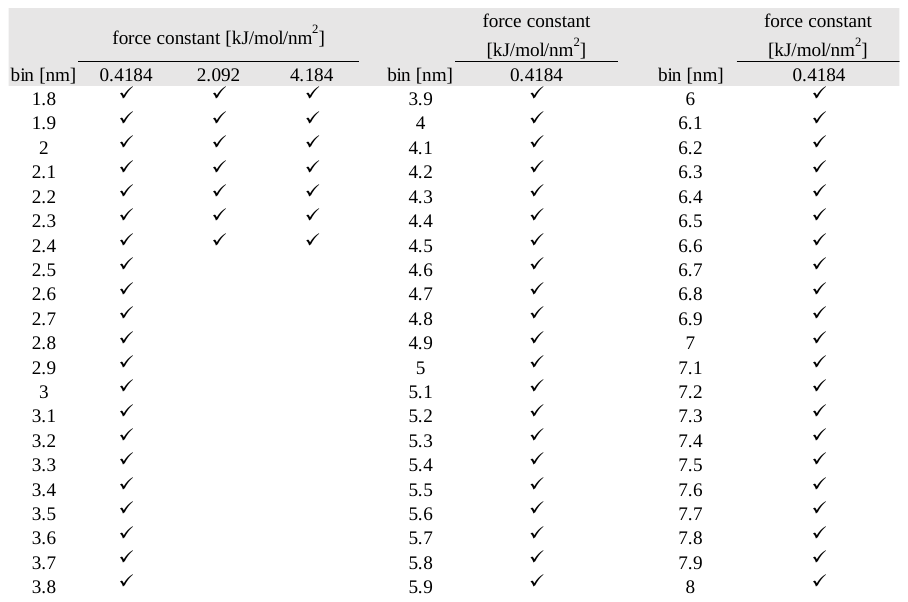

**Supporting Figures**

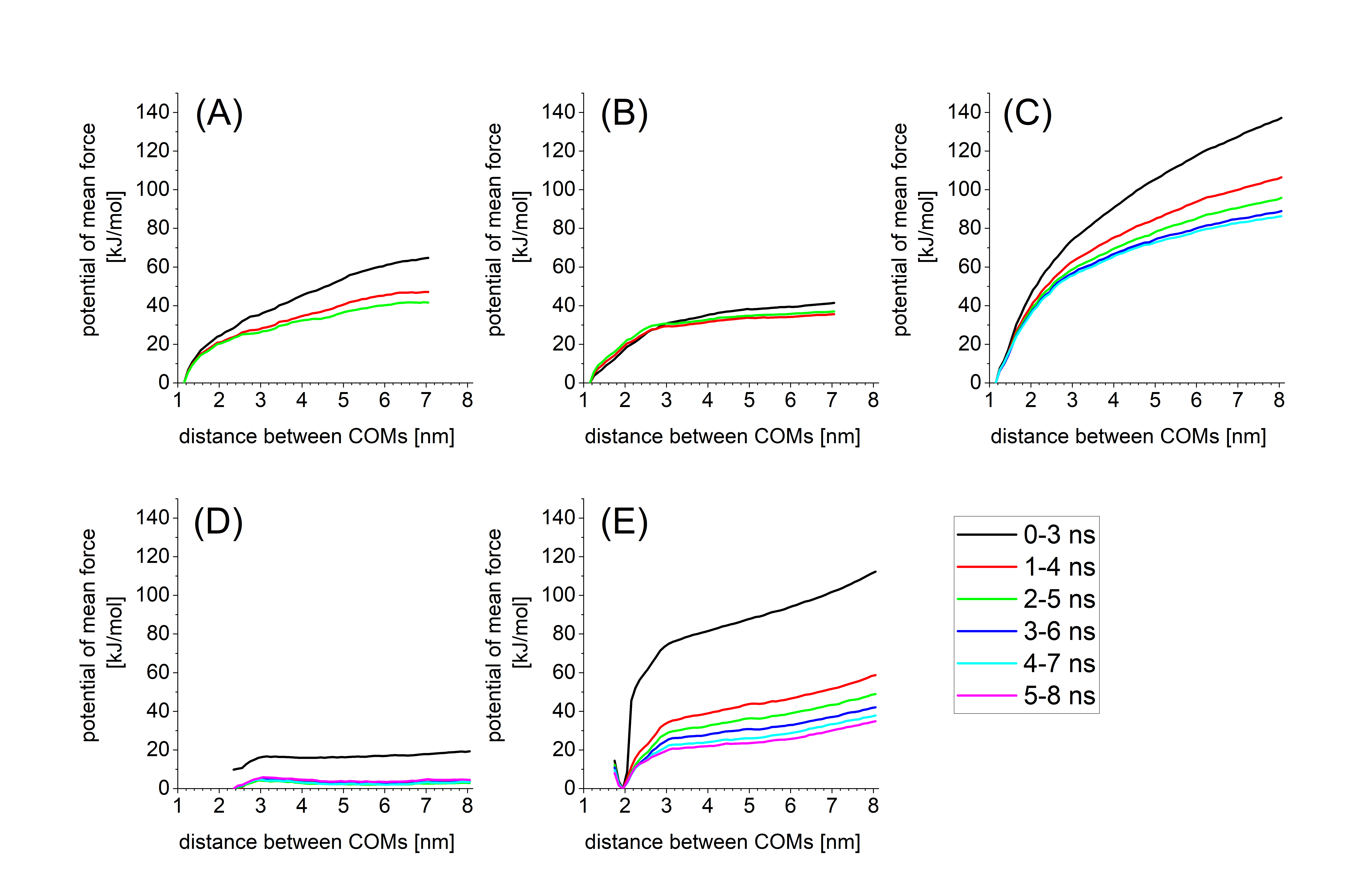

**Figure S1.** Potential of mean force calculation with shift of time domain. (A) S10 dissociation in Aβ_42_(M) system. (B) S10 dissociation in Aβ_42_(P) system. (C) S10 dissociation in Aβ_42_(M|R) system. (D) S19 dissociation in Aβ_42_(M) system. (E) S19 dissociation in Aβ_42_(M|R) system.
